# SOD1 controls neutrophil oxidative burst and microbial killing

**DOI:** 10.1101/2024.12.20.629705

**Authors:** Volker Brinkmann, Christian Goosmann, Andre Meier, Thomas Majer, Alessandro Foti

## Abstract

Neutrophils are immune cells specialized in producing large amounts of reactive oxygen species (ROS) to kill microbes. However, the mechanisms by which these cells regulate the balance of different ROS species and mitigate oxidative stress remain unclear. Here, we demonstrate that superoxide dismutase 1 (SOD1) plays a crucial role in ROS formation and antimicrobial activity in neutrophils. Our findings reveal that SOD1 modulates the ratio of superoxide (O_2_^-^) to hydrogen peroxide (H_2_O_2_) during the ROS burst, thereby supporting myeloperoxidase (MPO) enzymatic activity. By employing biochemical, cell biological, and genetic approaches, we show that SOD1 is crucial for ROS formation during NETosis and microbial infections, as it reduces oxidative stress and enables complete neutrophil activation. Impairment of SOD1 activity increases cysteine oxidation and lipid peroxidation. Neutrophils isolated from a patient with a SOD1 mutation exhibit decreased ROS production and impaired neutrophil extracellular trap (NET) formation. Our findings suggest that SOD1 is a novel regulatory factor in the oxidative burst that enables the full immunological response of neutrophils.

## Introduction

Neutrophils are the most abundant immune cells in human blood, playing a pivotal role in immune defense by sensing, capturing, eliminating invading microbes and communicating with other cells (1). Patients with neutropenia or with mutations affecting neutrophil function are highly susceptible to life-threatening infections (2). Neutrophils eliminate pathogens through three primary mechanisms: phagocytosis, degranulation, and the formation of neutrophil extracellular traps (NETs). Central to these processes is the generation of reactive oxygen species (ROS), a phenomenon known as the oxidative burst (3). This multistep process requires metabolic reprogramming, with increased glycolysis and activation of the pentose phosphate pathway (PPP) to supply NADPH (4). NADPH oxidase 2 (NOX2) transfers electrons from NADPH to molecular oxygen, producing superoxide (O_2_^-^), the precursor of most ROS. As both a mild reductant and oxidant, superoxide (O₂^-^) primarily reacts with itself, forming hydrogen peroxide (H₂O₂) in the presence of two protons (5). In most cells, H_2_O_2_ is the endpoint of the ROS cascade. However, neutrophils express high levels of myeloperoxidase (MPO), which uses H_2_O_2_ and halides to generate hypochlorous acid (HOCl), commonly known as bleach (6). Notably, neutrophils produce significantly larger quantities of reactive species than other cell types, which are critical for infection control. Consequently, patients with genetic disorders affecting ROS formation in neutrophils, such as glucose-6-phosphate dehydrogenase deficiency, NADPH oxidase 2 deficiency (chronic granulomatous disease, CGD), or myeloperoxidase deficiency, are prone to recurrent infections and autoinflammatory diseases (7).

ROS are intermediates of oxygen metabolism and are central for aerobic life. Most cells generate ROS as byproducts of their metabolism. However, if uncontrolled, ROS are toxic to cells (8). To mitigate oxidative damage, cells have evolved highly conserved systems spanning from archaea to humans. These include low-molecular-weight antioxidants, such as glutathione, and ROS-scavenging enzymes, such as catalase and peroxiredoxins. These systems maintain the “eu-oxidative stress”: the right amount of ROS in the appropriate locations (8). Neutrophils are specialized ROS producers, however, the redox mechanisms regulating immune processes in these cells remain poorly understood (9, 10). Neutrophils generate most ROS intracellularly in phagosomes and extracellularly at the plasma membrane. Plasma and phagosomal membranes are crucial to contain most ROS and prevent damage of cellular structures (11, 12). Moreover, redox-regulating enzymes, like glutathione peroxidase and thioredoxin are responsible for the appropriate functioning of neutrophils as antimicrobial and immune regulators (13–16).

To investigate how neutrophils balance ROS production during immune responses, we inhibited specific redox-regulatory pathways and tested oxidative burst, antimicrobial activity, and NET formation. We identified superoxide dismutase 1 (SOD1) as a crucial regulator of the oxidative burst. SOD1 is a highly conserved enzyme that catalyzes the dismutation of O_2_^-^ to H_2_O_2_. Mammalian cells express three distinct SODs, each localized to a specific compartment: SOD1 in the cytosol, SOD2 in mitochondria, and SOD3 in the extracellular space. SOD1- deficient neutrophils display impaired ROS burst, NETs formation, and killing of both *Candida albicans* and *Staphylococcus aureus*. Furthermore, lipid peroxidation and oxidative stress are enhanced in SOD1-inhibited neutrophils. Importantly, neutrophils isolated from a patient suffering of Amyotrophic Lateral Sclerosis (ALS) and carrying the mutation SOD1^R112Q^, also displayed reduced oxidative burst and NETs formation. Both pharmacological inhibition and loss-of-function mutations show that SOD1 is required to regulate the levels of O_2_^-^ during the oxidative burst, affecting H_2_O_2_ and HOCl formation in neutrophils.

## Results

### SOD1 enzymatic activity is required for NET formation

The oxidative burst is crucial for NET formation (17). To identify ROS pathways influencing NETosis, we tested 22 different compounds (Supplementary Table 1) that inhibit or modulate ROS-regulating enzymes and oxidative mechanisms in primary human neutrophils isolated from fresh peripheral blood of healthy donors. We stimulated the cells with the mitogen phorbol myristate acetate (PMA) and measured extracellular DNA release using the cell-impermeable dye SYTOX green as a proxy for NET formation. Compared to a vehicle control, three compounds, ATN-224, LCS1, and dithioethylcarbamate (DETC), inhibited extracellular DNA release by 50%, 70% and 80%, respectively. (Fig.1A). Interestingly, these three inhibitors all target SOD1 (18–20). The SOD1 inhibitors reduced NETs almost as effectively as the NOX2 and MPO inhibitors, diphenyliodonium (DPI) and 4-aminobenzoic acid hydrazide (4-ABAH), respectively (Fig. 1A). In addition, PX12 and PD1146176 — targeting thioredoxin-1 (Trx-1) and 15-lipoxygenase (15-LOX), respectively — achieved approximately 50% inhibition of DNA release (Fig. 1A). To confirm that DNA release in our assay represents NET formation, we stimulated neutrophils with PMA in the presence of the three SOD1 inhibitors and visualized NETs using the NET-specific antibody 3D9 (21) by immunofluorescence. The SOD1 inhibitors significantly reduced NET formation, consistent with the results of the SYTOX Green assay (Fig. 1C). To confirm these results with a more physiologically-relevant stimulus, we infected neutrophils with live *Candida albicans*, a human pathogen known to induce NETs (22). SOD1 inhibition reduced NET formation from 30% in vehicle-treated controls to 20%, 8%, and 12% when using ATN-244, LCS1, and DETC, respectively (Fig. 1D). As a positive control, DPI reduced NET formation to 10% and 5% following PMA and *C. albicans* stimulation, respectively (Fig. 1B, 1D). Furthermore, stimulation with PMA in the presence of exogenous recombinant SOD1 accelerated NETosis. Two hours after stimulation, NET formation reached 80% in the presence of exogenous SOD1 compared to only 20% without it (Fig. 1E). In contrast, adding exogenous catalase — which efficiently degrades ROS produced by SOD1 — or catalase together with SOD1, decreased NET formation (Fig. 1E). These results demonstrate that SOD1 activity promotes NET release in response to both mitogen and microbial stimulation.

**Figure 1.**
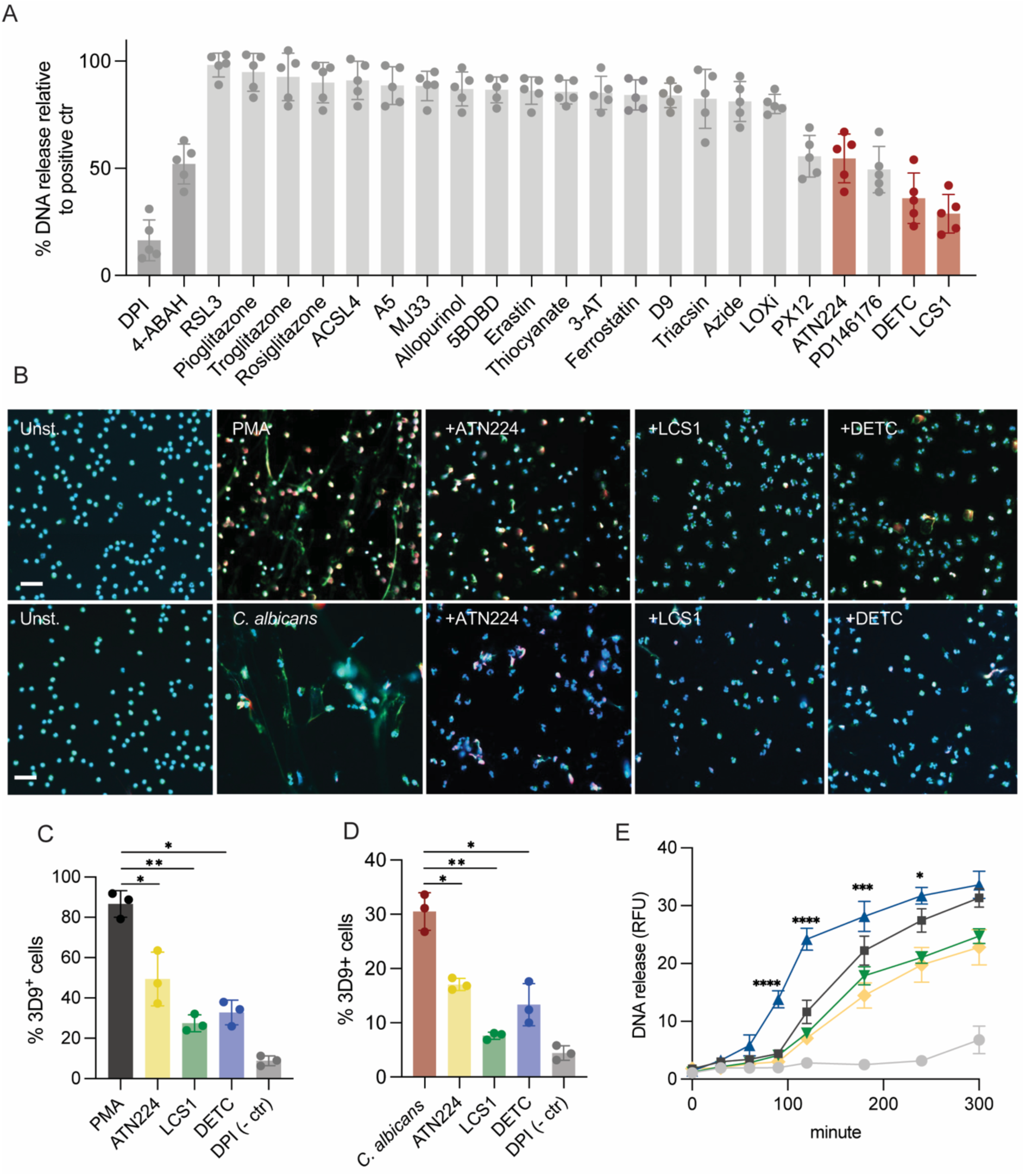
SOD1 inhibition blocks NETs formation. (A) Extracellular DNA release from human neutrophils stimulated with PMA and pretreated with inhibitors or modulators of redox pathways. Mean ± SEM of five independent experiments. Controls (dark gray), SOD1 inhibitors (red). (B) Staining of NETs with the 3D9 antibody. Human neutrophils were stimulated with PMA and *C. albicans* (MOI 1:1) in presence of SOD1 inhibitors (10 μM). Bars correspond to 10 μm. (C-D) 3D9 quantification of NETs. Human neutrophils stimulated with PMA or *C. albicans* in presence of SOD1 inhibitors (10 μM). Mean ± SEM of three independent experiments. *P &lt; 0.05, **P &lt; 0.01, unpaired T-test, two-tailed P value test. (E) NETs formation in the presence of exogenous recombinant human SOD1: naïve control (gray), PMA (black), exogenous SOD1 (blue) (10 μg/ml), exogenous catalase (green) (10 μg/ml) and exogenous SOD1+catalase (yellow). RFU, relative fluorescence unit. Mean ± SEM of three independent experiments. Statistical test one-way ANOVA and Bonferroni’s multiple comparisons test, **** &lt;0.0001, *** &lt;0.001, * &lt;0.05.

### SOD1 controls the level of ROS in neutrophils

To determine the specific effect of SOD1 activity during the ROS cascade, we inhibited the enzyme and measured different ROS species. First, we infected neutrophils with *C. albicans* and measured total ROS by the luminol assay, which detects all ROS species intra- and extra- cellularly. Surprisingly, SOD1 inhibitors, especially LCS1 and DETC, decreased the total ROS (Fig.2A). Then, we measured separately O_2_^-^, H_2_O_2_ and HOCl by using cytochrome C, Amplex red and aminophenylfluorescein (APF), respectively, upon stimulation with *C. albicans*. Neutrophils produced higher O_2_^-^ after *C. albicans* infection in the presence of SOD1 inhibitors, particularly with LCS1 and DETC (Fig.2B). Interestingly, H_2_O_2_ levels were reduced in presence of the SOD1 inhibitors, especially with LCS1 and DETC (Fig.2C). Consistently with the H_2_O_2_ results, HOCl was reduced when we inhibited SOD1 (Fig.2D). We repeated this assay using PMA as a ROS stimulus, to rule out any effects of *C. albicans* metabolic activity. Confirming our results with *C. albicans*, the SOD1 inhibitors also increased O_2_^-^ and reduced H_2_O_2_ and HOCl formation after PMA stimulation (S.Fig.2A-D). To control that the SOD1 inhibitors did not scavenge H_2_O_2_ or HOCl, we tested cell-free enzymatic MPO activity in the presence of the SOD1 inhibitors. The result showed no significant difference of MPO activity in presence of the three SOD1 inhibitors (S.Fig.2E). Together, these data indicate that SOD1 regulates the levels of different ROS after the activation of the oxidative burst. Finally, we showed that neutrophils treated with ATN-224, LCS1, and DETC displayed decreased antimicrobial activity against *C. albicans*, when compared to controls. Furthermore, we used a constitutively GFP expressing (GFP^+^) *C. albicans*, to monitor yeast growth for 6 hours in presence of neutrophils or medium alone as a control. In presence of SOD1 inhibitors, neutrophils displayed impaired *C. albicans* growth containment (Fig.2E). These data suggest that dysregulation of O_2_^-^ reduces the ROS burst and impairs the neutrophil antimicrobial capacity.

**Figure 2.**
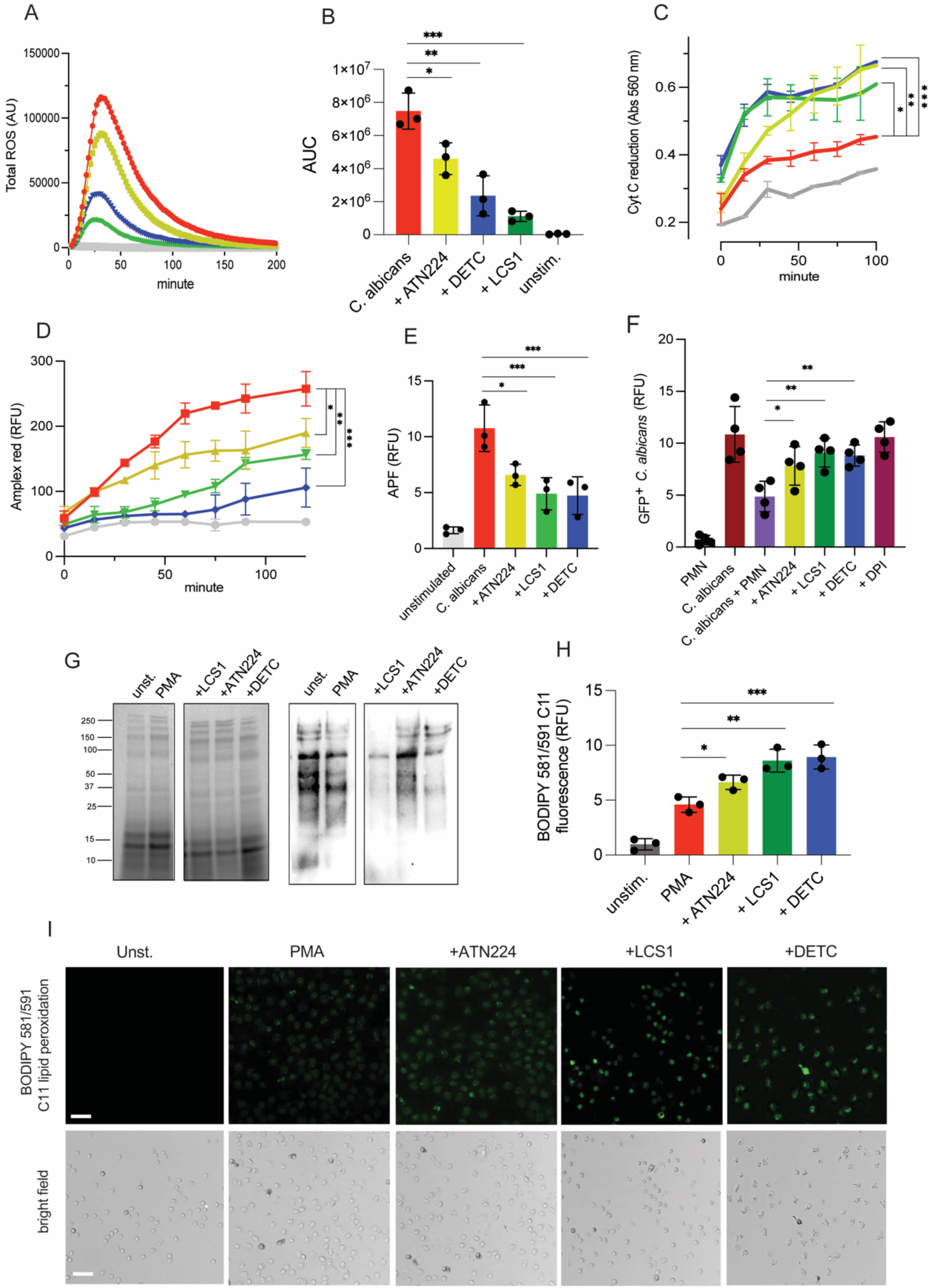
SOD1 controls the levels of ROS in neutrophils. (A-B) Human neutrophils infected with *C. albicans* (MOI 1:1) were tested for total ROS formation by luminol in presence of SOD1 inhibitors. (C) Superoxide was quantified by cytochrome C reduction assay. (D) Amplex red and (E) aminophenylfluorescein (APF) quantification of H2O2 and HOCl, respectively. (F) *C. albicans* growth (5 h) in the presence of human neutrophils pretreated with SOD1 inhibitors or controls. (G) Anti-thiol (cysteinyl) antibody-based quantification of oxidative stress on human neutrophil pretreated with SOD1 inhibitors and stimulated with PMA. SDS-PAGE (left) and Western Blot (right). (H-I ) BODIPY 581/591 C11 Lipid peroxidation quantification and live imaging of human neutrophils pretreated with SOD1 inhibitors and stimulated with PMA (1h). Bars correspond to 10 μm. (A, C and D) *C. albicans* (red) or *C. albicans* plus ATN224 (yellow), DETC (blue) and LCS1 (green). Statistical test one-way ANOVA with Bonferroni’s multiple comparisons test, **** &lt;0.0001, *** &lt;0.001, ** &lt;0.01, * &lt;0.05.

### Pharmacological inhibition of SOD1 leads to oxidative stress

We hypothesized that uncontrolled O_2_^-^ generation may damage neutrophils, limiting their functionality. Uncontrolled redox conditions and cysteines oxidation leads to dysregulation of the cellular functions (8). Additionally, lipid peroxidation is an oxidative damage occurring in case of unbalanced ROS formation, leading to ferroptosis and cell death (23). To test our hypothesis, we quantified the redox status of the whole neutrophil proteins and lipids. To analyze the redox status of the proteins, neutrophils were pretreated with SOD1 inhibitors, stimulated with PMA and lysed in the presence of Iodoacetyl Tandem Mass Tag (iodoTMT), an isobaric labelling reagent which binds to sulfhydryl-containing proteins for the detection of reduced cysteines. Thus, in case of oxidative stress, the probe would display decreased binding to cysteine thiols. We speculated that, if SOD1 inhibition leads to altered ROS formation, we would observe higher cysteine oxidation and fewer reduced thiols. Indeed, reduced thiols were less abundant in cells pretreated with SOD1 inhibitors and stimulated with PMA, as shown by Western Blot of cell lysates stained with an anti-TMT antibody (Fig.2F). This result indicates an oxidative imbalance in the absence of SOD1 activity. To test the effects of uncontrolled ROS on lipids, the BODIPY 581/591 C11 sensor was used to measure levels of lipid peroxidation. This sensor intercalates within the lipids of the cellular membranes and fluoresces after peroxidation. We found higher lipid peroxidation in neutrophils pretreated with SOD1 inhibitors compared to controls, in particular with LCS1 and DETC (Fig.2G). Together, these data demonstrate that SOD1 activity is required to regulate the redox state during the oxidative burst and allows optimal cellular activity.

### Intracellular localization of SOD1 in neutrophils

Due to their intrinsic chemical properties, ROS regulate biological pathways by proximity effects (24). In fact, the cellular localization of redox-regulatory enzymes is crucial to control the levels of oxidants. Hence, we determined the subcellular localization of SOD1 in naïve as well as in neutrophils stimulated with PMA, infected with *C. albicans* or *S. aureus* by immunofluorescence microscopy. Surprisingly, SOD1 was distributed in punctae, suggesting cytosolic aggregate-like structures (Fig.3A, S.Fig.3A-D). Interestingly, SOD1 and neutrophil elastase (NE), which localizes to primary granules, displayed a separated localization in naïve cells, however, they partially colocalized after PMA stimulation (Fig.3A). In infected neutrophils, SOD1 staining remained punctate, but localized around the phagolysosomes. We observed SOD1 in the cytosol as well as within membrane-bound vesicles by Transmission Electron Microscopy (TEM) (Fig.3B). Notably, SOD1 was partially detected within the phagosome, in neutrophils infected with *C. albicans* or *E. coli* (S.Fig.3E-H). As expected, MPO, a known primary granule marker, was also found within the phagosome and granules (S.Fig.3E-H). Altogether, our microscopy data indicate that SOD1 localizes to cytosolic aggregates, partially associated with membrane-bound vesicles, and locates within the phagosome during infection.

**Figure 3.**
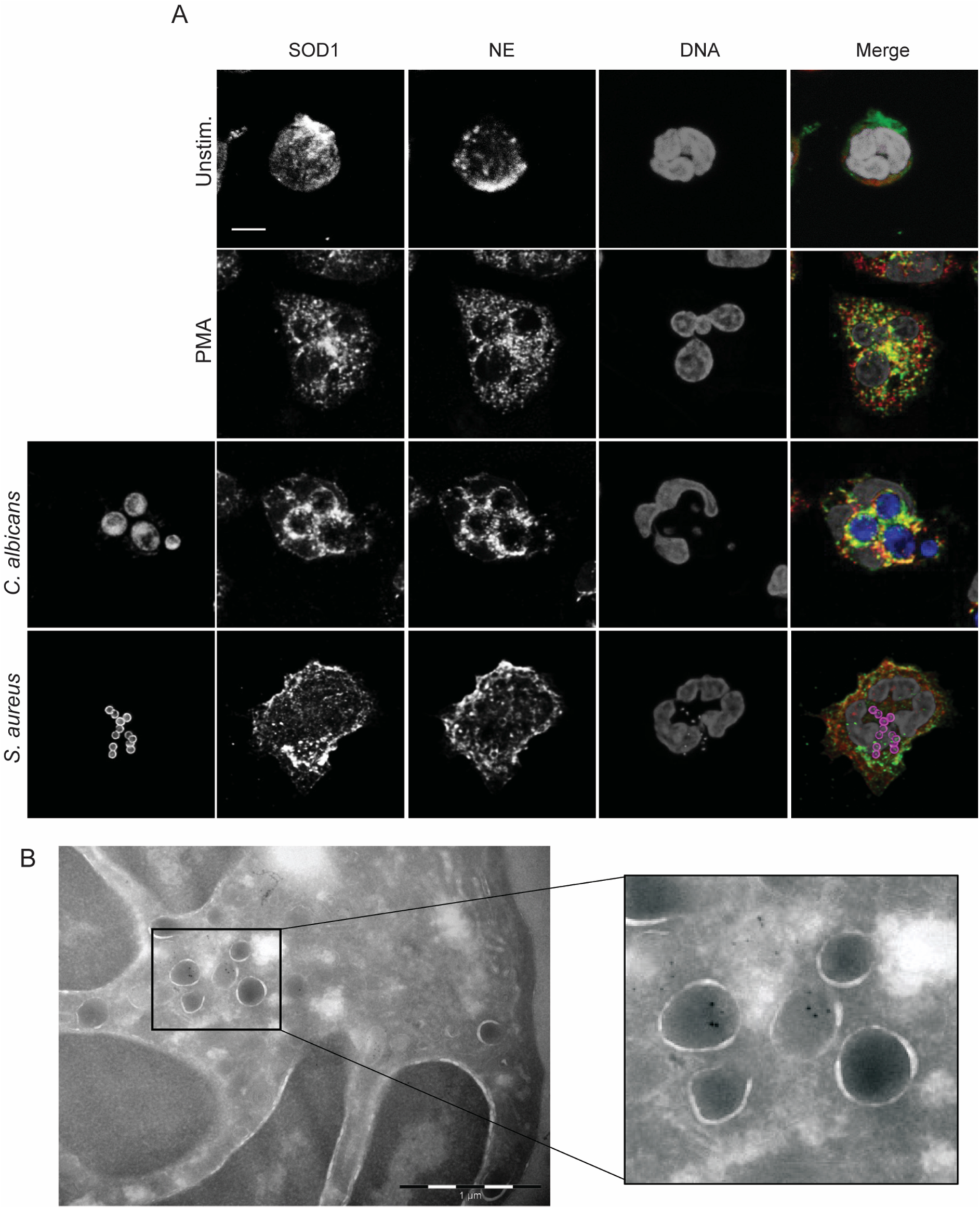
Intracellular localization of SOD1 in neutrophils. (A) Representative immunofluorescence images of human neutrophils stained with antibodies against NE (red), SOD1 (green), *C. albicans* (blue) or *S. aureus* (magenta). (B) Transmission electron microscopy of naïve human neutrophils. Large gold nano-particles indicate SOD1 and small gold nano-particles indicate MPO. Bars correspond to 2.5 μm (A) and 1 μm (B).

### SOD1^-/-^ mouse neutrophils are defective in ROS production, NET formation and containment of *C. albicans* growth

To obtain genetic evidence for the role of SOD1 in neutrophils, we moved to a SOD1-deficient mouse model (SOD1^-/-^). SOD1^-/-^ mice are smaller in size, age faster, display increased inflammation and susceptibility to infections compared to wild-type (WT) mice (25). Consistent with our inhibitor studies, SOD1^-/-^ peritoneal-elicited neutrophils made significantly less total ROS after *C. albicans* or PMA stimulation (Fig.4A-B). SOD1^-/-^ neutrophils also made fewer NETs than WT in response to *C. albicans* as determined with cell permeable (SYTO green) and impermeable (SYTOX orange) dyes after 5 hours (Fig. 4C). Interestingly, SOD1^-/-^ neutrophils challenged with *C. albicans* became permeable (SYTOX orange positive) but did not release NETs (Fig.4C-D; S.Fig.4). Around 40% of neutrophils, either WT or SOD1^-/-^, were SYTOX orange-permeable after *C. albicans* infection, as fluorometrically quantified (Fig.4E). However, quantification of the NET area showed that the SOD1^-/-^ neutrophils DNA fibers expand less than WT (Fig.4F). Moreover, we showed that SOD1^-/-^ neutrophils are less effective at reducing *C. albicans*^GFP+^ growth than WT (Fig.4G). Finally, we showed that SOD1^-/-^ neutrophils are less effective at killing *S. aureus* than WT cells by counting colony-forming units (CFU) (Fig.4H). As control, DPI-treated neutrophils, both WT and SOD1^-/-^, showed similar CFU counts. These results, consistent with our inhibitors data, demonstrate the critical role of SOD1 in ROS production after neutrophil activation and its impact on the antimicrobial capacity of these cells.

**Figure 4.**
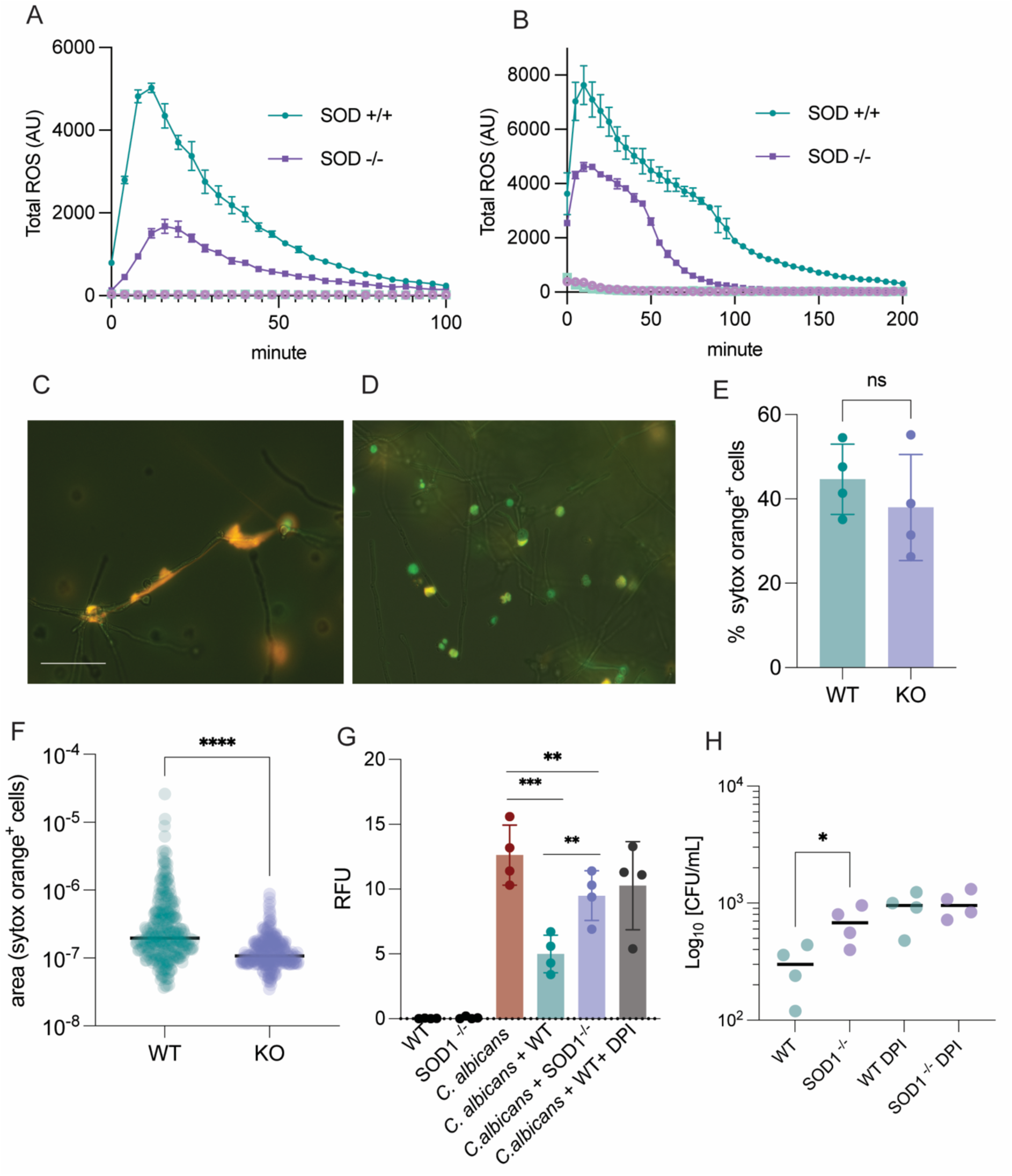
SOD1^-/-^ mouse neutrophils are defective in ROS production, NET formation, containment of *C. albicans* growth and killing of *S. aureus*. (A-B) Luminol assay of mouse neutrophils WT and SOD1^-/-^ stimulated with *C. albicans* (A) and PMA (B). Mean ± SEM of three independent experiments. (C-D) Live images of mouse neutrophils WT (C) and SOD1^-/-^ (D) after stimulation with *C. albicans* and stained for extracellular DNA (orange) and intracellular DNA (green) (5h). (E-F) Cell death quantification by SYTOX orange positive cells and quantification of extracellular DNA release by SYTOX orange area (inches^2^). (G) *C. albicans*^GFP+^ growth after co-incubated with neutrophils WT and SOD1^-/-^. (H) CFU counts one hour after infection of neutrophils WT and SOD1^-/-^ with *S. aureus* (MOI 1:5). Mean ± SEM of four independent experiments. (E-F) *P &lt; 0.05, **P &lt; 0.01, ***P &lt; 0.001, ****P &lt; 0.0001 unpaired T-test, two-tailed P value. (G-H) Statistical test one-way ANOVA with Bonferroni’s multiple comparisons test, *** &lt;0.001, ** &lt;0.01, * &lt;0.05. Bar corresponds to 50 μm.

### Human SOD1^R112Q^ neutrophils produce less ROS and NETs

Amyotrophic Lateral Sclerosis (ALS) is a neurodegenerative multifactorial disease that affects patients with a progressive loss of both upper and lower motor neurons that control voluntary muscles. Mutations on the *Sod1* gene are found in 12% of ALS cases (26). We obtained blood sample from an ALS patient carrying the variant SOD1^R112Q^ (27). The mutation R112Q generates insoluble SOD1 misfolded aggregates in motor neurons and affects mitochondria morphology and cellular metabolism (28). This mutation is proposed to impact the enzymatic activity of the protein (28). Mature human SOD1 is dimeric and arginine 112 lies at the dimerization face of the two monomers (Fig.5A). We isolated neutrophils from the ALS patient and characterized their ability to generate ROS and form NETs. Neutrophils from the ALS patient expressed an increased amount of SOD1 compared to cells from the healthy donor, as shown by Western Blot (Fig. 5C). The cellular distribution of SOD1 was similar in both ALS and healthy donor, as shown by immunofluorescence (S.Fig.5A). ALS neutrophils generated more O_2_^-^ in response to PMA than cells from healthy control (Fig.5B), but lower level of total ROS (Fig.5D). Moreover, ALS neutrophils showed a significant decrease and delay in NET formation over time, as shown by live microscopy and SYTOX green analysis (Fig.5E-F). Taken together, this case study recapitulated the pharmacological inhibition with the three different SOD1 inhibitors, and reinforces the role of SOD1 as a crucial regulator of the ROS burst and NET release in human neutrophils.

**Figure 5.**
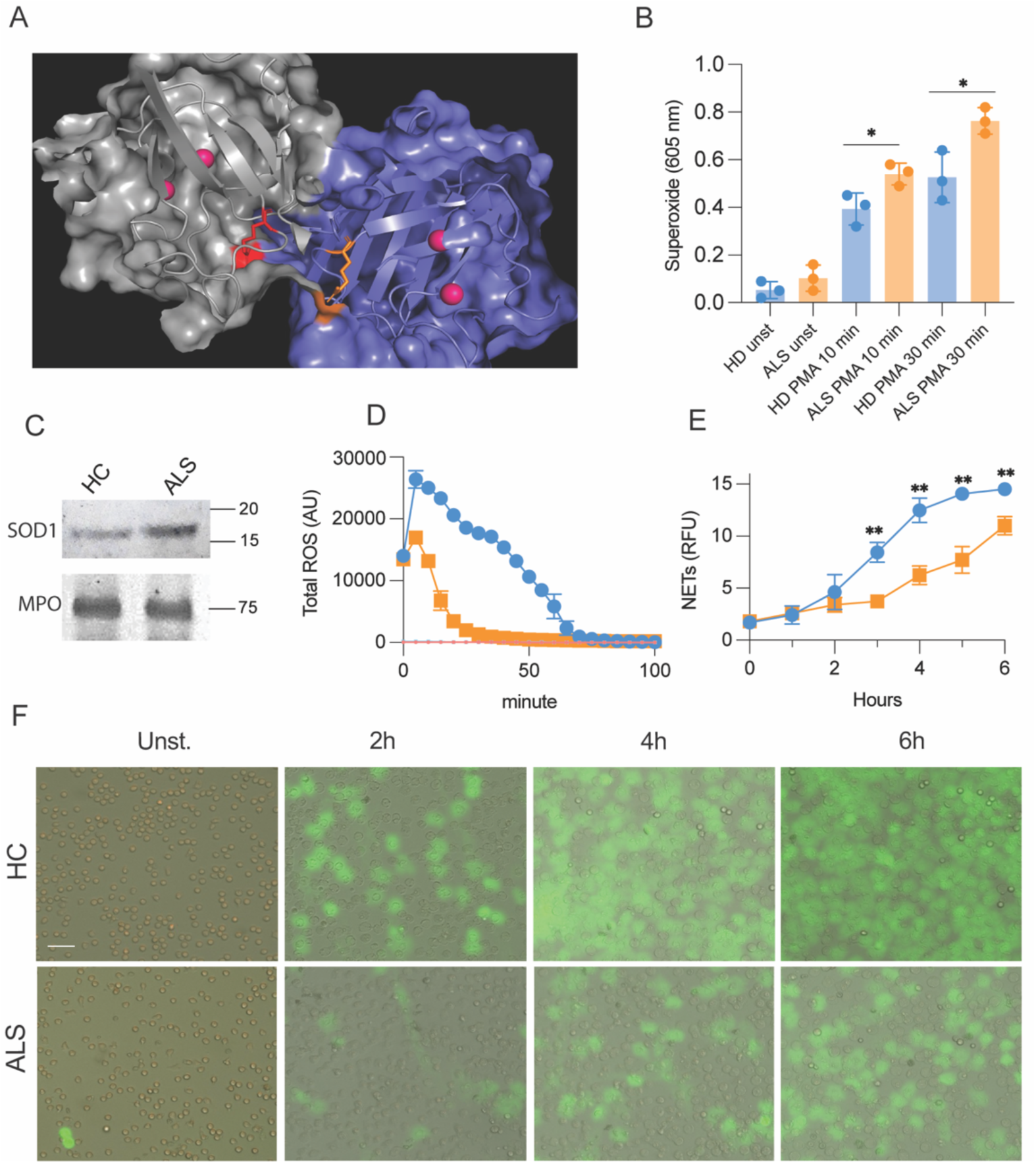
Human neutrophils SOD1^R112Q^ make less ROS and NETs. (A) Simulated 3D structure of SOD1 (PDB: 2C9V) and location of R112 obtained by PyMOL Molecular Graphics System. (B) Superoxide formation in healthy donor and ALS neutrophils after PMA stimulation measured by NitroBlue Tetrazolium (NBT) assay. *P &lt; 0.05, unpaired T-test, two-tailed P value. (C) Total ROS in ALS and healthy donor neutrophils after PMA stimulation. Healthy donor (blue) and ALS patient (orange). (D) Quantification and dynamic of NETs formation in ALS neutrophils; Healthy donor (blue) and ALS patient (orange). (E) Epifluorescence live images of NETs formation in healthy and ALS neutrophils after PMA stimulation. *P &lt; 0.05, **P &lt; 0.01, one-way ANOVA and Bonferroni’s comparisons test. Bar corresponds to 10 μm.

## Discussion

Activated neutrophils generate millimolar concentrations of ROS in few seconds. The activation of NOX2 produces O_2_^-^ and in few minutes depletes almost entirely the intracellular NADPH pool (6). This system is essential for neutrophils to combat microbes, control infections, and form NETs. Genetic impairments in ROS production render neutrophils ineffective, leading to recurrent infections (29). Here, to query neutrophil redox-regulation and NET formation, we screened a small-molecule library, which indicated SOD1 as novel regulator of the ROS burst and neutrophil activation.

SOD1 is a copper/zinc-dependent metalloenzyme initially identified as a ROS scavenger protecting cells from oxidative damage. Recently, numerous studies have indicated the regulatory role of SOD1 in a broad range of biological mechanisms, ranging from metabolism (30), to detoxification (31), inflammation (32) and RNA stabilization (33). However, its role in neutrophil biology remains underexplored. Given that neutrophils are “professional” ROS producers, we examined how SOD1 influences the oxidative burst. Notably, SOD1 converts O_2_^-^ to H_2_O_2_, a key substrate for MPO, which is critical for neutrophil antimicrobial activity. Our data show that SOD1 regulates ROS balance during the oxidative burst, supporting neutrophil antimicrobial responses. Pharmacological inhibition of SOD1 significantly reduces the ROS burst, underscoring its crucial role in neutrophil activation. To dissect the role of SOD1 in neutrophils, we used three inhibitors: LCS1, ATN224, and DETC. Inhibition of SOD1 increased O_2_^-^ levels while reducing H_2_O_2_ and HOCl production. LCS1 and DETC had stronger effects than ATN224, likely due to differences in molecular structure and inhibitory mechanisms. DETC and ATN224 specifically chelate zinc and copper ions, respectively, at SOD1’s active site, disrupting enzymatic activity (19, 20). LCS1 directly binds and inhibits SOD1 without altering its metal cofactors (18). Interestingly, exogenous recombinant SOD1 administered to stimulated neutrophils increased NETs formation. These data are consistent with previous observation from Palmer at al., (34), where exogenous SOD1 added to PMA-stimulated neutrophils enhanced NET formation. Genetic validation using SOD1-deficient mouse neutrophils confirmed the role of SOD1 in the ROS burst, critical for controlling *C. albicans* growth and *S. aureus* killing.

The absence of SOD1 activity altered ROS homeostasis in neutrophils, leading to oxidative stress. Uncontrolled oxidation damages biomolecules and disrupts cellular functions, causing protein aggregation via disulfide bridges (35) or ferroptotic cell death (23). Our data reveal that SOD1 deficiency in neutrophils increases cysteine oxidation and lipid peroxidation, hallmarks of oxidative stress. These findings support a model in which SOD1 activity maintains redox balance during neutrophil activation while sustaining the antimicrobial response.

The subcellular localization of redox-regulatory enzymes is critical for metabolic homeostasis. Mammalian cells express three superoxide dismutase isoforms: cytosolic (SOD1), mitochondrial (SOD2), and extracellular (SOD3). SOD1 is primarily cytosolic but also localizes to mitochondria (36). Electron microscopy and biochemical data from the Borregaard group showed that SOD3 locates within intracellular vesicles in neutrophils (37). *In vitro* infections using human macrophages, showed that SOD2 can be loaded into mitochondria-derived vesicle which fuse with the phagosome and promote killing of microbes (38). Our immunofluorescence studies show that SOD1 locates in a punctate cytosolic pattern. Electron microscopy shows that SOD1 is cytosolic as well as partially located into intracellular membrane-bound vesicle and within intraphagosomal space, in *C. albicans, S. aureus* and *E. coli* infections. These results suggest that SOD1 could actively contribute to the intraphagosomal ROS equilibrium. Further studies will be necessary to clarify the impact of SOD1 on intra- versus extra-cellular ROS dynamics and its impact in immune cells communication. Ultimately, we performed a case-study using neutrophils isolated from an ALS patient carrying the variant SOD1^R112Q^. ALS patients, in the later phase of their disease, suffer from several infections, mainly pneumonia (39). However, the overall risk of infection has not yet been linked to specific genetic profiles in ALS patients. SOD1^R112Q^ neutrophils displayed decreased total ROS, increased levels of O_2_^-^ and reduced NETs formation, confirming our previous pharmacological and genetic studies. Furthermore, our data show an increased superoxide formation in neutrophils after activation, which suggest possible alterations in cellular functionalities and immunological response.

Altogether, we propose a novel key step of the ROS burst pathway in neutrophils, which includes SOD1 as crucial regulator of the oxidative burst. We demonstrate that SOD1 activity is relevant in antimicrobial response against *C. albicans* and *S. aureus.* The discovery of a central role for SOD1 in the most ROS-producing immune cell opens a new avenue for studying the regulation of neutrophil antimicrobial activity and the consequences of oxidative stress in the innate immune system. This is not only important for basic innate immunology but may also provide key insights relevant to patients with metabolic pathologies, with further possible medical implications on patients carrying defective SOD1 mutations.

## Materials and Methods

### Chemicals

All reagents were purchased from common vendors of laboratory reagents such as Sigma Aldrich, Thermo Fischer Scientific or VWR Deutschland unless otherwise stated.

### Isolation of human neutrophils

The ethics council of the Charité Berlin (Germany) approved blood sampling and all donors gave informed consent according to the Declaration of Helsinki. The ALS patient consented to the use of biosamples as part of the registry study MND-NET (ethics committee number 19/12). Human neutrophils were isolated by a two-step density separation as described before (40). Briefly, freshly isolated peripheral blood was layered on an equal volume of histopaque 1119 and centrifuged at 800 g for 20 min. PBMCs and neutrophil layers were collected separately, washed with PBS with 0.2 % HSA and pelleted at 300 g 10 min. The neutrophil pellet was resuspended in PBS with 0.2 % HSA, layered on a discontinuous Percoll gradient (85 %-65 % in 2ml layers) and centrifuged at 800 g for 20 min. The neutrophil containing band was collected, washed in PBS with 0.2 % HSA and pelleted for 10 min at 300 g. The cell number was determined using a CASY cell counter (OMNI Life Science).

### Isolation of mouse neutrophils

Mouse breeding and isolation of peritoneal neutrophils were approved by the Berlin state authority Landesamt für Gesundheit und Soziales. SOD1-deficient (B6.129S7-Sod1^tm1Leb/J^) mice on a C57BL/6 background were obtained from Jackson Laboratories. Animals were bred at the Max Planck Institute for Infection Biology. Mice were housed in specific pathogen–free conditions, maintained on a 12-hour light/12-hour dark cycle, and fed ad libitum. C57BL/6 WT or SOD1-deficient mice 10-12 weeks old were injected i.p. with 1 ml of 7 % Casein solution in the evening and again after 12 h in the next morning. Three hours after the second injection, mice were sacrificed by rapid cervical dislocation. Murine neutrophils were isolated as previously described (41).

### Bacteria and yeast culture

S. aureus strain USA300 was cultured by inoculating 5 mL of sterile 3% (w/v) tryptic soy broth with a single colony and incubating at 37 C for 18 h with shaking. Bacteria were harvested, washed, and then resuspended in PBS. Bacteria were passaged into fresh TSB and incubated for 2h at 37 C at 180 rpm until the culture reached mid-logarithmic phase. Bacteria were harvested by centrifugation for 5 min at 2000 x g, washed twice with RPMI 1640 medium supplemented with 5mM HEPES pH7.4 and adjusted to OD600 = 0.1. In experiments, the cultures were used at a final OD600 of 0.05, which equated to roughly 50 x 10^6^/mL. *E. coli* XL1-Blue (Stratagene) was cultured overnight at 37°C in LB plus tetracycline (22).

*C. albicans* (clinical isolate SC 5314) and *C. albicans*^GFP+^ (42) were maintained on yeast peptone dextrose (YPD) agar plates. A single colony of *C. albicans* was inoculated in 5 ml YPD broth for 18 h at 37 C. From this intermediate culture, 0.1 ml was removed and added to 4.9 ml of fresh YPD broth that was incubated at 37 C for an additional 4 h (43). *C. albicans* cells were washed twice by centrifugation in PBS, quantified by absorbance at 600 nm and confirmed by hemocytometer counting. Detailed information about *C. albicans*^GFP+^ strain generation are in Gerami-Nejad et al., (42).

### Microbial killing assay

Rate of bactericidal activity of neutrophils was determined and calculated by a one-step bactericidal assay (44). Freshly isolated 10^6^ neutrophils were incubated with 5×10^6^ opsonized *S. aureus* in 1 ml of RPMI with 0.1% human serum albumin, with final ratio of bacteria to neutrophils of 5:1. Samples were continuously rotated and incubated at 37 C and after 45 min were placed at 4 C. Neutrophils and phagocytosed bacteria were pelleted by centrifugation at 100 x g for 10 min. Supernatants were plated on TSA plates to determine the number of bacteria that were not phagocytosed. The pellets containing neutrophils and phagocytosed bacteria were washed twice with PBS and then hypotonically lysed using water at pH 11.0. After lysis, samples were pelleted by centrifugation at 300 x g for 10 min to remove the neutrophils debris and the supernatant containing the bacteria was plated on TSA plates.

For the candicidal assay, *C. albicans*^GFP+^ was coincubated with human or mouse neutrophils (10_6_) at MOI 1:1 for 5 h in RPMI medium at 37 C. After 5 h, the growth of *C. albicans*^GFP+^ was fluorometrically quantified by GFP emission, using Fluoroskan Ascent fluorescence spectrometer, and analyzed by epifluorescence microscopy. *C. albicans* ^GFP+^ growth was validated by correlating GFP fluorescence to metabolic activity measured by tetrazolium salt (XTT) reduction assay (45).

### ROS quantification

#### Total ROS

To assess total ROS production, 10^5^ neutrophils were activated with 50 nM PMA or *C. albicans* (MOI 1:1), after inhibitors treatment or vehicle. ROS production was measured by monitoring luminol (50 μM) luminescence in the presence of 1.2U/ml horseradish peroxidase (17).

#### Superoxide

The cytochrome c reduction assay was performed as previously described (46). Briefly, cells were suspended in 200 µl RPMI containing 75 µM ferricytochrome c and were activated by adding PMA or *C. albicans* (MOI 1:1). The absorbance at 550 nm with a 490 nm reference filter was on a Versa Max microplate reader.

Alternatively, superoxide was measured by the colorimetric NBT assay (47). 10^5^ neutrophils were incubated in RPMI containing NBT and simulated with PMA for 60 min. As negative controls, some cells were incubated in NBT solution containing PMA with 5 mM DPI. After incubation, cells were washed twice with PBS, then once with methanol, and air-dried. The NBT deposited inside the cells were then dissolved, first by adding 120 mL of 2 M KOH to solubilize cell membranes and then by adding 140 mL of DMSO to dissolve blue formazan with gentle shaking for 10 min at room temperature. The dissolved NBT solution was then transferred to a 96-well plate and absorbance was read on a microplate reader at 620 nm.

#### Hydrogen peroxide

The Amplex Red-dependent fluorescence assay was performed according to the manufacturer’s instructions (Thermo Fisher) and previously published methods (48).

Amplex red (final concentration 5 µM) was added to the neutrophils resuspended in RPMI medium in a 96-well plate. For studies evaluating the effect of specific SOD1 inhibitors on H_2_O_2_ formation, neutrophils were incubated for 15 min at 37 C with the different inhibitors. All reactions were adjusted to a final volume of 100 µl with RPMI. The fluorescence was monitored using a Fluoroskan Ascent fluorescence spectrometer at an excitation wavelength of 530 nm and an emission wavelength of 590 nm. Background values, defined as the mean fluorescent values of Amplex Red reaction mixture diluted in RPMI, were subtracted from all readings.

#### Hypohalous acids

Neutrophils were seeded onto a glass-bottomed dish and loaded with APF (5 µM) by incubation for 30 min at room temperature. Dye-loaded cells were stimulated with PMA (50 nM) or *C. albicans* (MOI 1:1). Hydrogen peroxide as positive control was used at the concentration of 500 µM, to stimulate hypohalous acids formation in neutrophils. Fluorescence images were acquired twice in each experiment (before and 10 min after the stimulation with PMA) using a Leica SP8 confocal microscope. The excitation wavelength was 488 nm, and the emission was filtered using a 505–550 nm barrier filter.

### NETs detection and quantification

Purified human neutrophils were seeded at a density of 10^5^ cells in 96-well plates for measurements of NET formation by SYTOX Green. For immunofluorescence stainings and live imaging, neutrophils were seeded in µSlide 8 Well ibiTreat dishes. Cells were treated with inhibitors 30 min before induction of NET formation. Cells were stimulated with PMA (100 nM) or *C. albicans* (MOI 1:1).

Mouse neutrophils were seeded at 10^5^ in 24-well plates in RPMI (Gibco) containing penicillin/streptomycin (Gibco) and glutamine (Gibco), 1% murine DNase -/- serum, and murine granulocyte colony-stimulating factor (100 ng/ml; PeproTech). Forty-five minutes after seeding, cells were stimulated with live *C. albicans* for 5 hours. NETs were quantified with microscopic images after staining unfixed cells with the cell-permeable DNA dye SYTO green and the cell-impermeable DNA dye SYTOX Orange.. The area of DNA-positive regions was quantified using the software ImageJ following automatic default thresholding, and total pixel area was converted to inches squared (inch²) using image calibration. Watershed method was applied for segmentation. For immunofluorescence staining, neutrophils were seeded at 10^5^ per well in µSlide 8 Well glass-bottom dishes (ibidi) and treated with *C. albicans* (MOI 1:1) for 5 hours. Cells were fixed with 2% paraformaldehyde (PFA) for 30 min at room temperature. After washing with phosphate-buffered saline (PBS), cells were permeabilized with 0.1% Triton X-100 at 4°C for 5 min, washed again with PBS, and blocked with 3% bovine serum albumin (BSA). The NETs specific antibody 3D9 was added to samples overnight at 4°C. After washing with PBS, secondary antibodies were added in 3% BSA for 30 min, and after washing with PBS, images were acquired on a Leica SP8 confocal microscope. DNA was counterstained with DAPI (1 µg/ml).

### Neutrophil lysate preparation

Cell lysates were prepared from stimulated or resting neutrophils, in presence or absence of SOD1 inhibitors. Cells were seeded in culture medium in 1.5 ml microcentrifuge tubes at 1×10^7^cells/ml. After spinning down for 10 min at 300 g, cells were resuspended in RIPA buffer (Thermo Fischer) with 20 μM neutrophil elastase inhibitor GW311616A (Biomol) and Halt protease inhibitor cocktail (PIC, Thermofisher Scientific), gently mixed and incubated at 37 °C with gentle rotation. Cells were gently mixed and centrifuged at 1000 g (30 s) to collect all residual liquid. Freshly boiled 5 X sample loading buffer (50 mM Tris-HCl pH 6.8, 2% [w/v] SDS, 10% glycerol, 0.1% [w/v] bromophenol blue, 100 mM DTT) was added to samples which were then briefly vortexed and boiled (98 °C) for 10 min with agitation and flash frozen in liquid nitrogen for storage at –80 °C.

### Thiol oxidation

Human neutrophils (5×10^5^ per condition) were stimulated with PMA after pretreatment with inhibitors or vehicle. Cells were lysed as mentioned above but without reducing agents. In addition, the lysis buffer contained iodoTMT labeling agent as indicated by the manufacturer (iodoTMTsixplex™ Isobaric Label Reagents, Thermo Fisher). After labeling, the lysates were analyzed by Western Blot and stained with and anti-TMT antibody (TMT Monoclonal Antibody 25D5, Invitrogen) to detect the level of reduced thiols.

### Lipid peroxidation

Human neutrophils naïve or stimulated with PMA, in presence of SOD1 inhibitors or vehicle, were measured for lipid peroxidation by using the Sensor BODIPY™ 581/591 C11 as indicated by the manufacturer (Thermo Fisher). Briefly, 10^5^ cells were seeded in a 96 well plate in RPMI in presence 2 µM of BODIPY 581/591 C11 for 30 min. After it, cells were washed two times with RPMI. Subsequently, cells were treated with inhibitors or vehicle for 30 min. After it, cells were washed with RPMI, resuspended in RPMI and PMA (50 nM). After one hour cells were examined by fluorescence microscopy using a Leica SP8 confocal microscope.

### Immunogold Electron Microscopy of ultrathin cryosections

Neutrophils were harvested, fixed in 2% PFA and 0.05% glutar-di-aldehyde, gelatin-embedded and infiltrated with 2.3 M sucrose according to the method described (49).

For TEM analysis, ultrathin sections were cut at -110°C with a RMC MTX/CRX cryo-ultramicrotome (Boeckeler Instruments Inc., Tucson AZ, USA) transferred to carbon- and pioloform-coated EM-grids and blocked with 0.3% BSA, 0.01M Glycin, 3% CWFG in PBS. The sections were incubated with appropriate dilutions in the same buffer of mouse monoclonal antibody directed against SOD1 and rabbit polyclonal antibody directed against MPO. Secondary antibody-incubations were carried out with donkey-anti-rabbit and donkey-anti-mouse antibodies coupled to 12 nm or 6 nm gold particles (Jackson Immuno Research,West Grove, PA, USA). Specimens were then contrasted and embedded with uranyl-acetate/methyl- cellulose following the method described (50) and analyzed in a Leo 906 transmission electron microscope (Zeiss, Oberkochen, Germany) operated at 100 kV. Images were recorded using a side-mount Morada digital camera (SIS, Münster, Germany)

### Immunofluorescence staining

Figure 1B: human neutrophils were seeded into 12 well chamber slides (ibidi) with removable silicone gaskets and treated as indicated. After incubation, cells were fixed with 2% PFA/PBS for 30 min, permeabilized with 0.5% Triton X-100 for 30 sec and blocked for 30 min (PBS + 3% normal donkey serum, 3% cold water fish gelatin, 1% BSA and 0.05% Tween 20). Cells were stained with 3D9 mouse mAb for 60 min (5 µg/ml in blocking buffer). Bound antibody was visualized with donkey anti mouse secondary antibodies coupled to AlexaFluor488. DNA was stained with Hoechst 33258. After removal of the gaskets, the slides were mounted using Mowiol, and whole-slide scans were prepared using a ZEISS AxioScan Z1 at 10x magnification. The percentage of 3D9-positive nuclei was determined using QuPath 5.0 software. Figure 3A: human neutrophils were seeded onto round coverslips (diameter 12 mm) in 24-well plates and treated as indicated. Cells were fixed and stained according to the protocol for Fig. 1b. Coverslips were mounted with Mowiol and analyzed on a Leica SP8 confocal microscope. Figure S.5A: neutrophils were resuspended at 10^6^ cells/ml in RPMI containing 0.02% human serum albumin and 10 mM Hepes, and 200 µl of cell suspension was dropped into µSlide 8 Well glass-bottom dishes (ibidi). After stimulation, a pH shift fix was performed [3% PFA–K-Pipes (80 mM) (pH 6.8) for 5 min followed by 3% PFA-borax (100 mM) for 10 min]. Cells were permeabilized using 0.1% Triton X-100 at 4°C for 2 min. Cells were then blocked (5% goat serum, 1% fish gelatin, 2% BSA) for 1 hour followed by incubation with primary antibody in 0.1% BSA at 4°C overnight, washed once, and then incubated with Alexa Fluor–coupled secondary antibodies (Life Technologies) at room temperature for 20 min. After three PBS washes and one incubation in distilled water for 20 min, cells were imaged in PBS using either Leica SP8 confocal microscopy or total internal reflection microscopy. DNA was counterstained with Hoechst (5 µg/ml) or DAPI (1 µg/ml).

### Live imaging of neutrophils

Cells were resuspended in RPMI (Gibco) supplemented with Hepes (5 mM) and 0.2% HSA supplemented with 500 nM SYTOX Green and 2.5 uM DR and seeded at a density of 5 × 10^5^ cells per well into µSlide 8 Well ibiTreat dishes (ibidi). After SOD1 inhibitors treatment and NET induction, images were collected at 10-min intervals on a EVOS epifluorescence imaging system. The recording of DR (cell-permeable DNA dye) was used to track individual nuclei over time and to determine nuclear area. In case of human neutrophils, the recording of SYTOX Green (cell-impermeable dye) was used to determine permeability of cells. In case of mouse neutrophils, cells were stained with the cell-permeable DNA dye SYTO Green and the cell-impermeable DNA dye SYTOX Orange.

## Acknowledgements

We thank Arturo Zychlinsky for advices on the project and discussing the manuscript. We thank C.J. Harbort, Dhiren Ferise Patel and Sarah Ruddle for reading the manuscript.

## Author Contributions

VB, CG and AF performed experiments and analyzed the data AM and TM cared for the patients

AF designed the study and wrote the manuscript

---

**Supplementary Figure 1.**
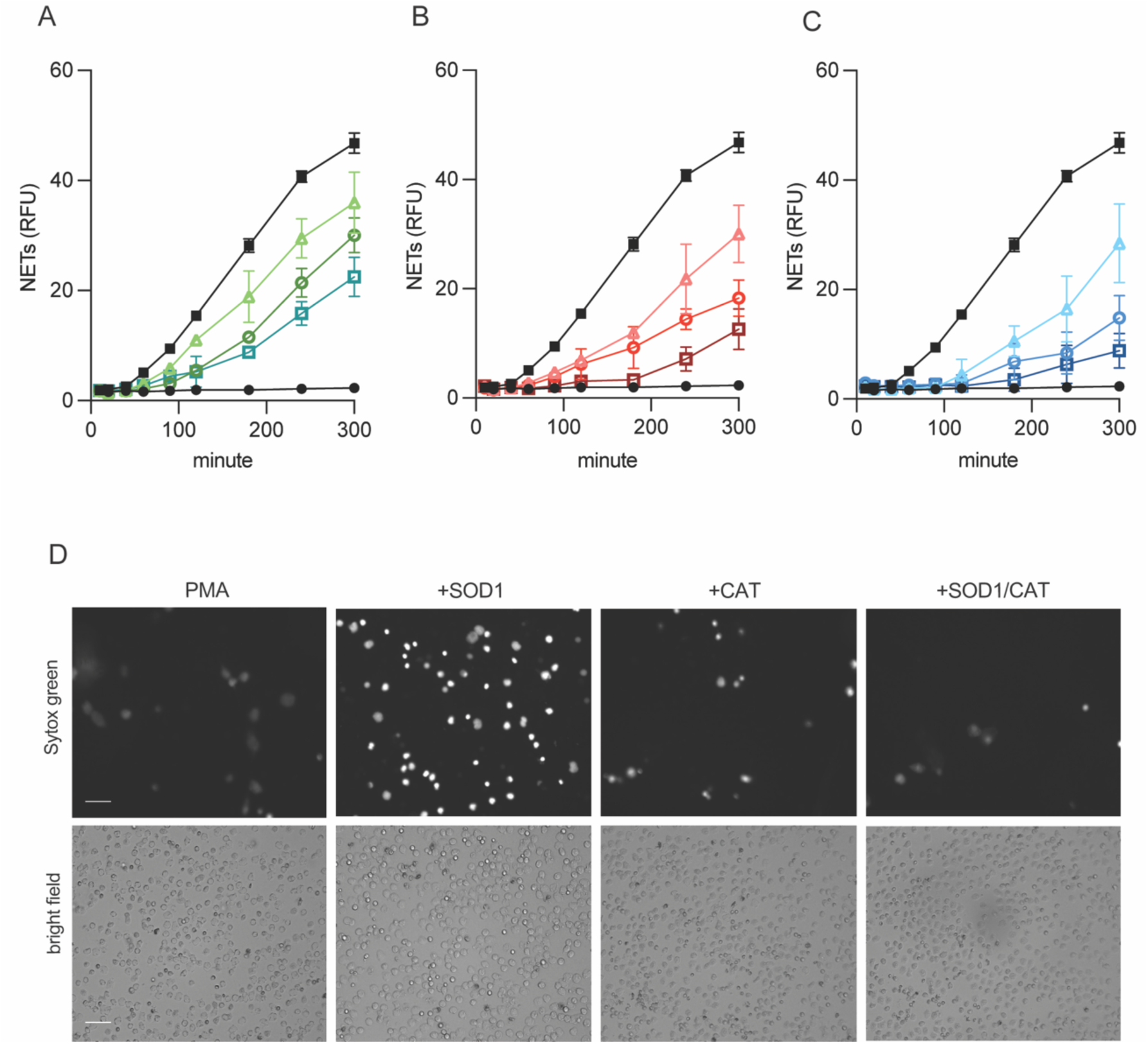
S**O**D1 **inhibition blocks NETs formation:** (A-C) NETs quantification, dynamic and dose response (1, 10 and 100 μM) of ANT224 (A), LCS1 (B) and DETC (C) inhibitors on human neutrophils stimulated with PMA (black) measured by SYTOX green assay. Mean ± SEM of five independent experiments. (D) Epifluorescence live images of DNA release after PMA stimulation (2h) in presence of exogenous SOD1, catalase and SOD1/catalase (10 μg/ml). Bars correspond to 10 μm.

**Supplementary Figure 2.**
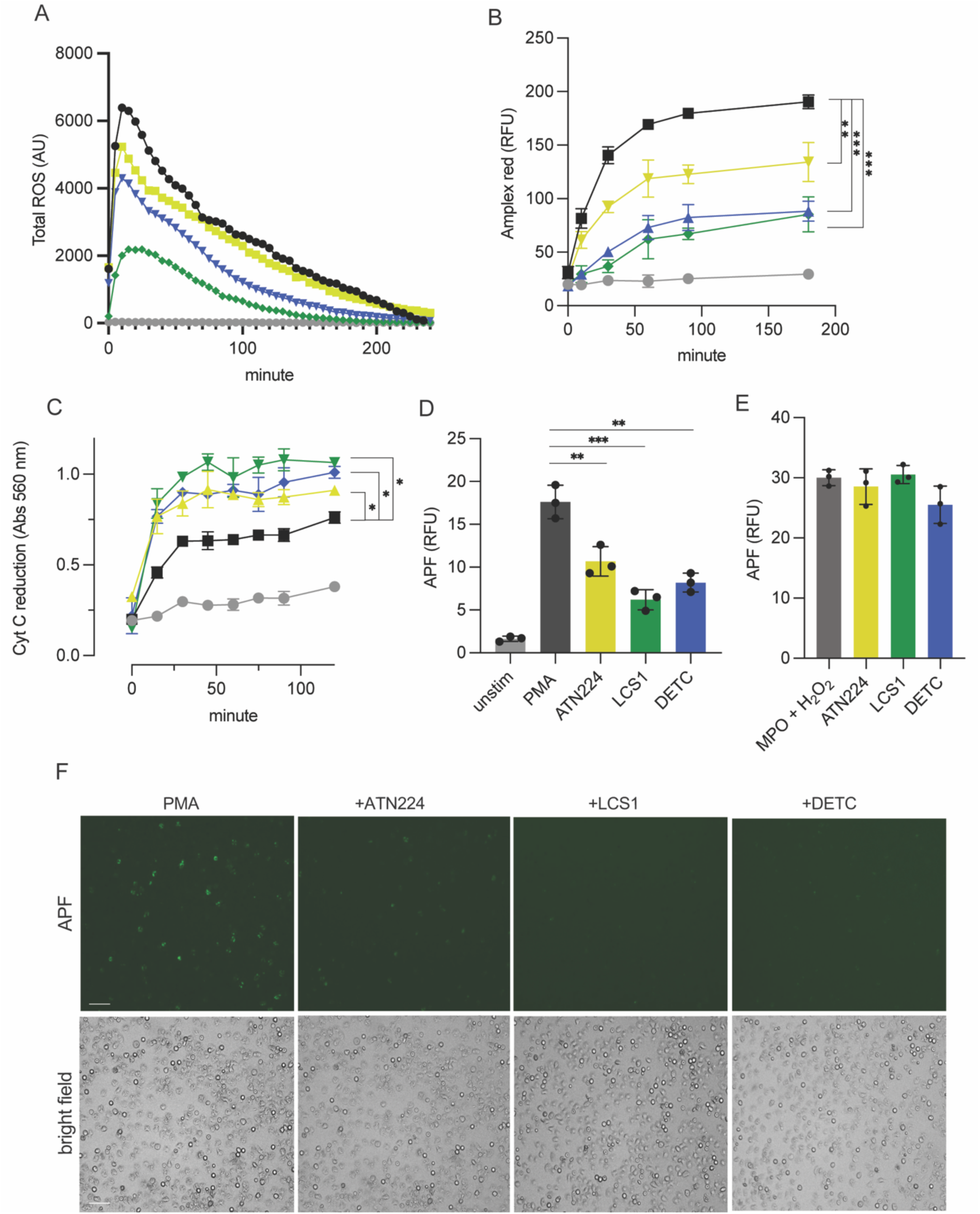
S**O**D1 **controls the levels of ROS in neutrophils:** (A) Luminol, (B) Amplex red and (C) Cytochrome C assays of human neutrophils stimulated with PMA in presence of SOD1 inhibitors. (D) Aminophenylfluorescein (HOCl sensor) quantification of human neutrophils stimulated with PMA and in presence of SOD1 inhibitors. (E) *In vitro* MPO activity in presence of SOD1 inhibitors. (F) Aminophenylfluorescein (HOCl sensor) live imaging of human neutrophils treated with SOD1 inhibitors and stimulated with PMA for 1 hour. Mean ± SEM of three independent experiments. (A-C) PMA (black) or PMA plus ATN224 (yellow), DETC (blue) and LCS1 (green). Statistical test one-way ANOVA with Bonferroni’s multiple comparisons test, *** &lt;0.001, ** &lt;0.01, * &lt;0.05.*P &lt; 0.05. Bars correspond to 10 μm.

**Supplementary Figure 3.**
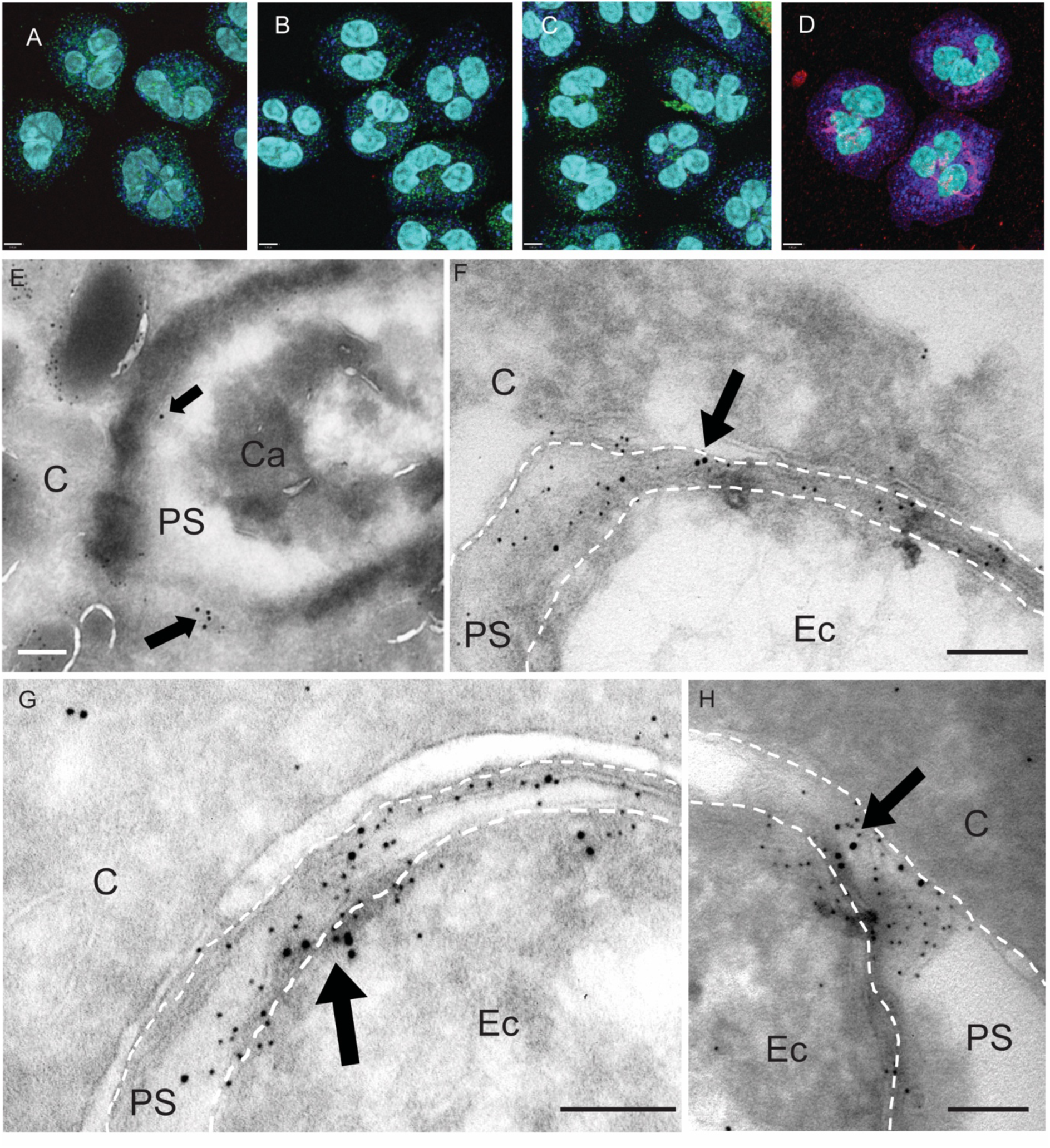
I**n**tracellular **localization of SOD1 in neutrophils:** (A- D) Confocal images of human neutrophils stained for MPO (blue), SOD1 (green in A-C, magenta in D) and DNA (light blue) with four different primary SOD1 antibodies. Transmission electron microscopy (TEM) of human neutrophils infected with *C. albicans* (E) or *E. coli* (F-H). Small and large gold nanoparticles indicate MPO and SOD1, respectively. Arrows indicate SOD1. Bars correspond to 1 μm (A-D) , 0.1 μM (E) and 250 nm (F-H). C, cytosol; PS, phagosomal space; Ca, *C. albicans*; Ec, *E. coli*.

**Supplementary Figure 4.**
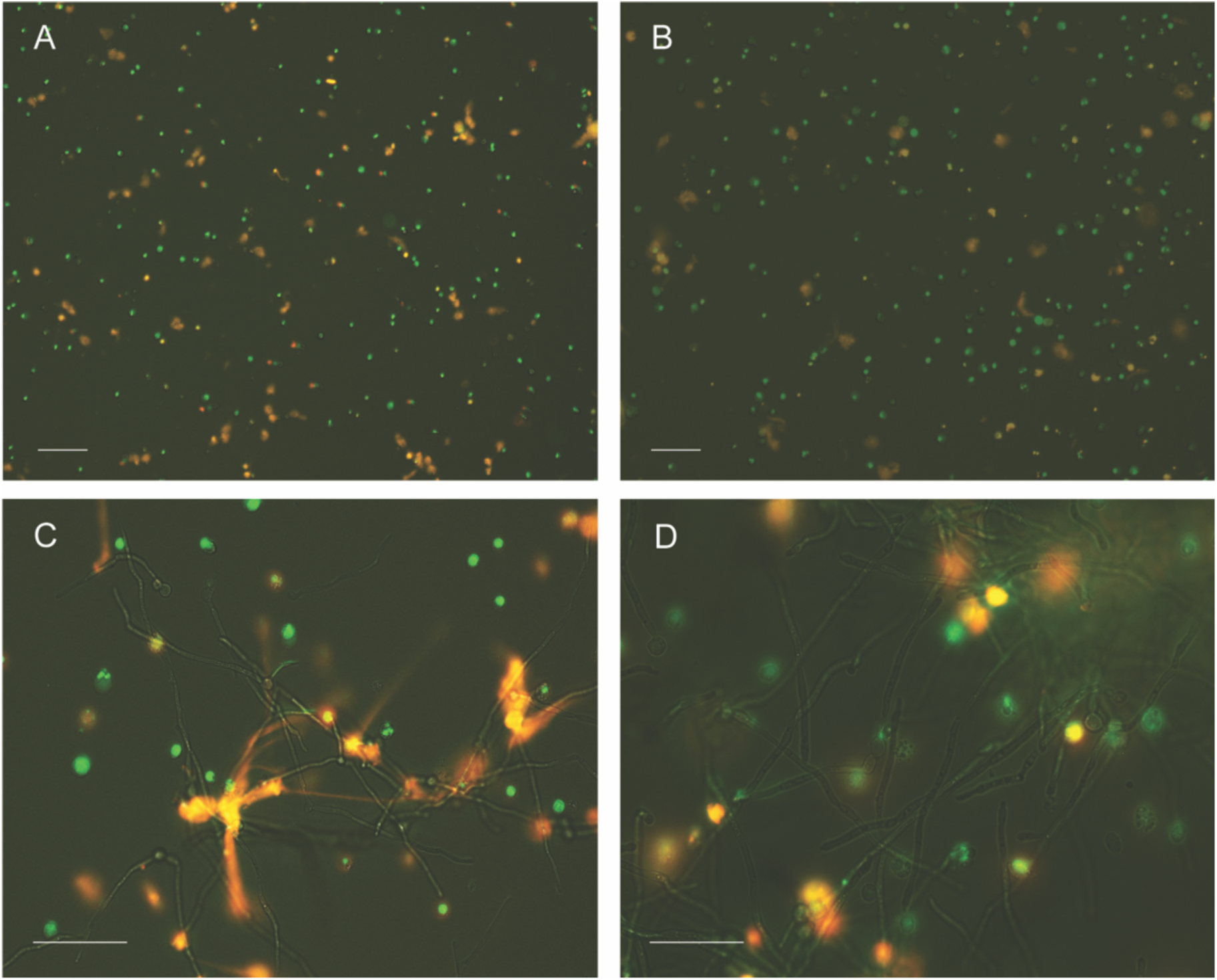
SOD1^-/-^ mouse neutrophils are defective in ROS production, NET formation, containment of *C. albicans* growth and killing of *S. aureus*: (A-B) Epifluorescence live images (5h) of naïve mouse neutrophils WT (A) and SOD1 ^-/-^ (B). (C-D) Mouse WT (C) or SOD1 ^-/-^ (D) neutrophils after infection with *C. albicans* (MOI 1:1). Neutrophils were stained with SYTOX orange and SYTO green for extracellular DNA (orange) and intracellular DNA (green), respectively. Bars correspond to 75 μm (A-B) or 50 μm (C-D).

**Supplementary Figure 5.**
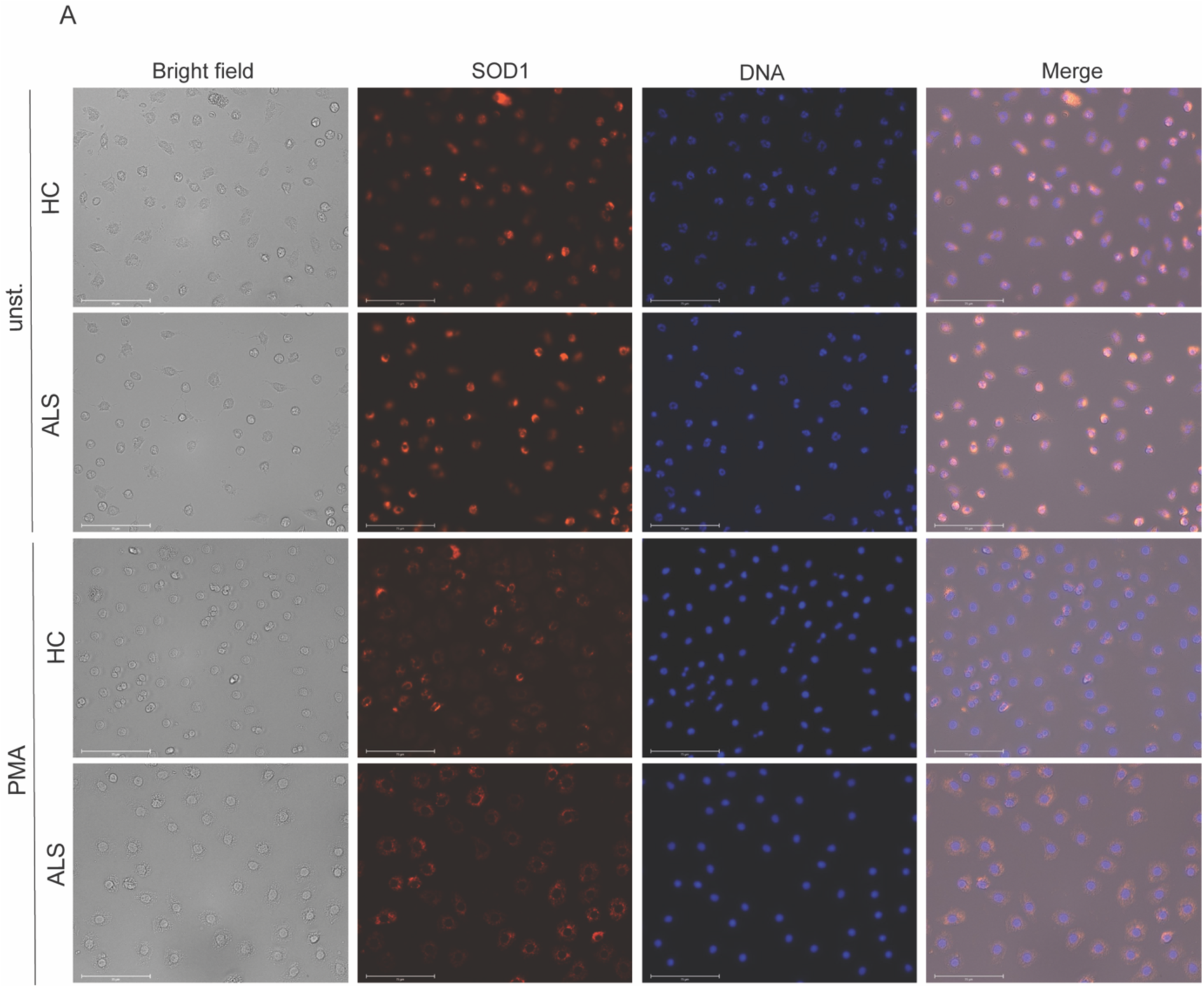
H**u**man **neutrophils SOD1^R112Q^ make less ROS and NETs:** (A) immunostaining of SOD1 in healthy donor and ALS patient neutrophils, naïve or stimulated with PMA (50 nM) for 1 h. DAPI (blue) and SOD1 (red). Bars correspond to 75 μm.

**Supplementary table 1:**

| **compound** | **target** | **concentration** | **reference** |
| --- | --- | --- | --- |
| Erastin | initiator of ferroptosis | 10 μM | Sun et al. 2020 |
| Ferrostatin | ferroptosis inhibitor | 20 μM | Miotto et al, 2020 |
| RSL3 | inhibitor of glutathione peroxidase 4 | 10 μM | Yu et al, 2019 |
| PX12 | thioredoxin-1 inhibitor | 5 μM | Guang-Zhen et al, 2015 |
| D9 | thioredoxin reductase inhibitor | 5 μM | Bradford et al, 2024 |
| MJ33 | peroxiredoxin VI inhibitor | 25 μM | Intae et al, 2013 |
| Triacsin | long fatty acyl CoA synthetase inhibitor | 10 μM | Dechadt et al, 2017 |
| LOXi | arachidonate 5-lipoxygenase inhibitor | 2 μM | Kahnt et al, 2022 |
| ACSL4i (PRGL493) | Acyl-CoA Synthetase Long Chain Family Member 4 inhibitor | 5 μM | Yu et al, 2023 |
| A5 | complex II inhibitor | 10 μM | Wojtovich et al, 2009 |
| Rosiglitazone | ferroptosis inhibitor | 5 μM | Syang et al, 2024 |
| Pioglitazone | agonist of PPARγ | 1 μM | Chenyang et al, 2022 |
| Troglitazone | ferroptosis inhibitor | 1 μM | Doll et al, 2017 |
| LCS1 | superoxide dismutase1 inhibitor | 10 μM | Somwar et al, 2011 |
| ATN224 | superoxide dismutase1 inhibitor | 10 μM | Lee et al, 2013 |
| DETC | superoxide dismutase1 inhibitor | 10 μM | Khazaei et al, 2009 |
| Thiocyanate | myeloperoxidase substrate, antioxidant | 100 μM | Manoharan et al, 2024 |
| Allopurinol | xanthine oxidase inhibitor | 10 μM | Sekine et al, 2023 |
| 3-AT | catalase inhibitor | 50 μM | Ueda et al, 2003 |
| 5BDBD | P2X4 receptor antagonist | 2 μM | Coddou et al, 2019 |
| PD146176 | 15-Lipoxygenase inhibitor | 5 μM | Bocan et al, 1998 |
| Azide | Mitochondrial electron transfer chain inhibitor | 50 μM | Eiermann et al, 2022 |
| 4-ABAH (- ctr) | myeloperoxidase inhibitor | 100 μM | Kenny et al, 2017 |
| DPI (- ctr) | NADPH oxidase 2 inhibitor | 100 nM | Piszczatowska et al, 2019 |

